# Structure-resolved fold annotation of the medicinal-leech salivary-EV cargo: a Ly-6/uPAR three-finger scaffold and a structurally dark fraction

**DOI:** 10.64898/2026.10.26.762066

**Authors:** Hongfen Cen, Fang Zhao, Juan Zhao, Xiangrong Tong, Zichao Liu, Feng Zhao

**Affiliations:** Yunnan Key Laboratory of Leech Bioresources and Industrialization, College of Agriculture & Life Sciences, Kunming University, Kunming 650214, China; Key Laboratory of Ethnic Medical Resources Research and Southeast Asian International Cooperation in Yunnan Province, School of Tea and Coffee, Pu’er University, 6 Xueyuan Road, Pu’er, Yunnan 665000, China; Yunnan International Joint Laboratory of Digital Conservation and Germplasm Innovation and Application of China-Laos Tea Resources, Pu’er University, 6 Xueyuan Road, Pu’er, Yunnan 665000, China

**Keywords:** medicinal leech, *Hirudinaria manillensis*, salivary extracellular vesicles, structural proteomics, protein fold annotation, Foldseek, Ly-6/uPAR three-finger fold, dark proteome

## Abstract

Many proteins that medicinal leeches package into salivary extracellular vesicles (EVs) carry no interpretable sequence annotation, so the least-characterised part of a pharmacologically important secretome lacks even a fold assignment. A companion Astral-DIA study of *Hirudinaria manillensis*^1^ nominated 17 ultra-enriched candidate EV cargo proteins. Here we ask whether structure prediction and structural search can assign folds where sequence methods fail, and which candidates remain unresolved under prespecified criteria. With an offline pipeline (ESMFold, TM-align, Foldseek against AFDB-SwissProt and PDB100, UniRef90 search, protein-to-genome alignment), the candidates separate into intracellular/membrane machinery and a secreted, cysteine-dense fraction of three proteins with no Pfam domain. The predicted structure of the leech Protein-sleepless homolog g21166 supports its assignment to the Ly-6/uPAR three-finger fold (TM 0.58 vs CD59; best significant PDB match Lynx1); no function is inferred. g436 and g20791 returned no match meeting the criterion (alnTM ≥ 0.5, E ≤ 0.05) in either database, g436 even after homolog-informed modelling. Gene structure and splice junctions support a conserved locus for the two-Trypsin-segment model g8114 (catalytic unit not established). A Boltz-2 thrombin pilot did not support target engagement, and a pre-registered nicotinic test was terminated as undecidable. The results yield a structure-informed shortlist for experimental follow-up.

## Introduction

Hematophagous leeches deploy a pharmacologically rich salivary secretome that underlies both their traditional medicinal use and modern anticoagulant drug development. Part of this secretome is carried by extracellular vesicles (EVs), and in the medicinal leech *H. manillensis* a companion comparative-proteomics study^1^ nominated 17 ultra-enriched candidate EV cargo proteins that do not correspond to canonical EV markers. Several of them, including eight “uncharacterised” proteins, carry no interpretable sequence-level annotation; that study therefore concluded that structure-based fold annotation was required to interpret them and recommended AlphaFold^2^/Foldseek^3^ as the next step.

Such sequence-dark proteins are the hardest part of a secretome to interpret. Without a recognised domain or an annotated homolog, and often without a place in an interaction network, they receive no functional hypothesis and are easily set aside, although they may include lineage-specific secreted proteins. Protein structure is more conserved than sequence, so predicted structures searched against structure databases can recover relationships that sequence comparison misses.^2,3^ For small, cysteine-rich proteins, however, single-sequence models are often of low confidence, and the absence of a structural match is easily over-read as a novel fold. Whether structure-based annotation can resolve such cargo therefore depends on separating a genuine fold assignment from a non-detection, under criteria fixed in advance.

Here we ask whether structure prediction and structural search can assign folds to the salivary-EV cargo that sequence methods leave unannotated, and which candidates remain structurally dark. Specifically, we test (i) which of the small candidates are predicted to fold into stable scaffolds whose cysteine spacing is consistent with the predicted disulfide pairings (no disulfide bond is measured here); (ii) whether the Protein-sleepless^4^ homolog adopts the three-finger fold^5^ implied by its sequence family; and (iii) whether any candidate lacks a structural match under a prespecified criterion and, if so, whether this reflects missing homolog information or the structure itself. Answering these questions turns part of the “uncharacterised” label into testable fold hypotheses and yields a structure-informed shortlist for experimental work. All structural statements are confidence-quantified predictions; a target-engagement layer was only partly executed and is reported as such (Section 2.5). All computation was performed offline; no candidate sequence or structure was submitted to any online server (Experimental Section).

### Relationship to the companion paper

The comparative Astral-DIA proteome — the two-compartment quantification (salivary-EV-enriched PM vs whole-body pharmacopoeial powder FDPMD), differential-abundance analysis, in-silico MISEV^6^/EV signature, within-family pharmacopoeia isoform divergence and the nomination of the 17 candidates — is the subject of a separate companion manuscript (Zhao et al., *PeerJ*, ms 144559, under review^1^). The present paper reports only the structural follow-up that study recommended. To avoid redundant publication, Section 2.1 recapitulates the proteomics only as the starting point, with citation; the compartment, DEP, GO and isoform analyses are not reproduced here.

### Experimental Section

#### Biological material and ethics

The proteomic resource re-analysed here was generated from adult *Hirudinaria manillensis* (synonym *Poecilobdella manillensis*) obtained from Yunnan Yuyu Biopharmaceutical Co. Ltd. (Kunming, China); no field collection was performed, and no collection permit was therefore required. *H. manillensis* is an invertebrate not subject to formal vertebrate-animal ethics review at the participating institution; the work nevertheless followed institutional best-practice guidelines for invertebrate welfare. No new animal material was used for the present study, which is computational throughout. Full husbandry and sampling details are given in the companion paper.^1^

#### Provenance and non-duplication

The proteome, differential-abundance analysis, MISEV/EV signature, subcellular partitioning, pharmacopoeia isoform divergence and 17-candidate nomination are reported in the companion manuscript^1^ (Astral-DIA, DIA-NN,^7^ 2,818 protein groups; 1,749 differential proteins in its corrected classification — 737 continuous BH-controlled differentially abundant proteins plus 1,012 presence/absence candidates). We reuse only the candidate accessions and their reported enrichment as the starting point; no differential-abundance result is re-derived or re-claimed here.

#### Candidate extraction & characterization

Candidate sequences were extracted by accession from the 25,347-entry *H. manillensis* protein database used in the companion study. Length, MW, pI, GRAVY, net charge, cysteine count/density and putative disulfide count were computed. Protein-domain annotation used HMMER^8^ 3.4 `hmmscan` vs a local Pfam-A^9^ (E<1e-5).

#### Structure prediction

ESMFold^10^ structure prediction was attempted for all 17 candidates (`facebook/esmfold_v1` via HuggingFace `transformers`^11^); the six candidates ≤260 aa were first folded on CPU and all 17 were then run in a single GPU batch whose per-candidate record — status, mean pLDDT, attention chunk size, peak allocation, wall time and input sequence SHA-256 — is reported in Table S1. Sixteen models were produced and all sixteen were searched; one candidate, g18761.t1 (3,570 aa), could not be folded (CUDA out-of-memory at every chunk size 64/32/16/8; peak allocation 89.57 GiB on a 96 GB device), so it is absent from every structural analysis in this paper and its exclusion is a hardware-capacity limit, not a selection. MSA-based high-confidence modelling (offline ColabFold/AlphaFold3) is the scoped extension.

#### Fold assignment & structural search

Pairwise TM-align^12^ (tmtools) against a reference fold panel (RCSB PDBs^13^); same-fold threshold TM≥0.5; both reference-and chain-normalized TM are reported. Foldseek 10 `easy-search` of models vs a local AFDB^14^-SwissProt^15^ database (`easy-search`, `-e 10`; output fields query, target, alntmscore, evalue, prob and theader); a *significant Foldseek match* is defined as a hit with alnTM ≥ 0.5 and E ≤ 0.05, and existence is evaluated over all reported hits for a query rather than over its top-scoring hit alone; a query with no such hit is reported as lacking a significant match. Note that foldseek’s `alntmscore` is normalized by min(alignment, query, target) length and is therefore not a full-length qTM/tTM, so a significant match is not by itself a same-fold assignment.

#### Homolog search and within-model MSA ablation

Candidate sequences were searched against a local UniRef90^16^ MMseqs2^17^ database (snapshot built on 2 August 2026; 121,389,642 entries) with `-s 5.7 --max-seqs 2000` using the same pipeline applied to all null-model objects; no sequence left the machine. For g436, two Boltz-2^18^ 2.2.1 arms were run that differed only in the MSA input (depth-103 A3M versus `msa: empty`), with identical parameters (`predict --override --num_workers 4 --no_kernels`) and seeds 101–103 per arm. Structural searches used the highest-confidence seed of each arm (with-MSA seed 103, no-MSA seed 101). Confidence comparisons are made within Boltz-2 only; ESMFold and Boltz-2 pLDDT are not compared across models. A PDB100-wide Foldseek search was also performed (August 2026; foldseek 10.941cd33; PDB100 snapshot of 1 January 2025; the same `-e 10`, additionally reporting qtmscore, ttmscore and bits). Note that foldseek applies its default prefilter and `--max-seqs 1000`, so this is a PDB100-wide search rather than an algorithmically exhaustive all-vs-all; “no significant match” is therefore reported as *under this search*, not as an absolute claim of a new fold.

#### Nicotinic target-engagement test (pre-registered, terminated)

A pre-registered AlphaFold3 interface test was designed for g21166 against an AChBP dimer as a soluble surrogate for the nicotinic receptor, with the interface statistic defined as the maximum chain-pair ipTM between ligand and either receptor chain, and the decision rule requiring the candidate to exceed every member of a fold-matched, non-nAChR null model. Compute prerequisites were met (AlphaFold3 v3.0.4 with the complete v3.0 databases, CRC32C-verified). The test was terminated as undecidable at the null-model construction stage. First, no length stratum contained candidate, null model and positive controls simultaneously: predicted mature lengths were 84 aa (g21166), 57-66 aa (snake three-finger toxin nulls) and 68-74 aa (positive controls); an alternative Ly-6/uPAR null pool (71-110 aa) matched the candidate but contained no positive control within +/-8 aa. Second, functional negativity could not be established from the available evidence: both candidate null families are themselves enriched for nicotinic-receptor ligands, so removing annotated binders yields a set for which activity is *unreported* rather than *excluded*. One further consideration bears on the scoring method itself. Interface confidence scores of the ipTM family have a documented failure mode in within-family discrimination: for the Kazal inhibitor domains, an inhibiting and a non-inhibiting domain of the same protein both score above the conventional 0.75 threshold, and an engineered two-substitution variant that experimentally loses inhibition is statistically indistinguishable from wild type.^19^ We therefore treat ipTM-family scores as exploratory descriptors rather than confirmatory discriminators absent a target-system-specific validation. We report no interface result for this target and retain the nicotinic assignment as a fold-and structural-similarity-based hypothesis.

#### Comparative gene structure and transcript support (g8114 only)

The g8114.t1 protein (1,313 aa) was aligned to three published chromosome-level hirudinean assemblies with miniprot^20^ 0.18-r281 (`-t 8 --gff`), the query being the g8114.t1 protein sequence (deposited with this paper): *Hirudinaria manillensis* (GCA_034509925.1, ASM3450992v1, isolate Hman_Guangxi — the assembly from which the query gene model derives), *Hirudo nipponia* (GCA_040113095.1, isolate rbyz1) and *Whitmania pigra* (GCA_041430665.1, isolate K615); local copies were retrieved 2026-07-04. Intron positions were expressed in query-protein coordinates from the `Target` attribute of each CDS feature and compared between species with a ±3-aa tolerance, chosen because the cross-species protein identity (0.51–0.52) makes the alignment imperfect at exon boundaries. Splice sites were read directly from each genome; GT–AG and the recognised minor GC–AG class were both counted as canonical. Because the same query protein and the same aligner were used for all three genomes, the three mappings are not methodologically independent; they test conservation of exon–intron architecture, not transcript identity. For transcript-level support, public *H. manillensis* RNA-seq runs were aligned to the same assembly with HISAT2^21^ 2.2.2 (`-p 32 --dta`, paired-end) and junction-spanning reads were counted at MAPQ ≥ 60 against the miniprot-predicted intron coordinates: three salivary-gland libraries (BioProject PRJNA861860) and four head libraries (SRR26151944 from PRJNA1019887; SRR26541746, SRR26541752 and SRR26541753 from PRJNA1032729). A prespecified admissibility rule required ≥ 100 uniquely mapped reads at the locus for a library to be interpretable; SRR26541752 (95 reads) and SRR26541746 (49) fell below it and were excluded before the junctions were examined, leaving two head libraries drawn from two separate studies. Detection was tested against expression-matched control introns on the same chromosome (n = 9,948) with a one-sided (lower-tail) binomial test. That test assumes introns are conditionally independent given expression, which is optimistic for introns of one transcript, and the controls are matched at gene level whereas the g8114 regions are intragenic, so the reported *P* values are approximate. No de-novo genome annotation was performed.

#### Interaction context

This study used the node and edge tables deposited with this paper to render the network with igraph^22^ and to query membership of the leads; no network was constructed here. The plotting script records these tables as a STRING^23^ network filtered at score ≥900; because we cannot verify that attribute against the tables as delivered, we refer to it throughout as the network used here rather than as a high-confidence STRING network. The plotting script records the origin of those tables as the vendor’s DIA report accompanying the proteomics service; the original report is not retained in this project, so that origin — and whether the tables were processed after delivery — has not been independently verified by us. Network parameters (STRING score ≥900) are likewise traceable only to those tables and that script annotation.

#### Target engagement (pilot executed; full layer incomplete)

The planned layer comprised human hemostasis and nicotinic-receptor targets from RCSB (thrombin 1PPB/4HTC, FXa 1FJS/2P16, FXIa 5TKC, plasmin 1BML, AChBP surrogate), complexes via AlphaFold3^24^-multimer/Boltz-2, ΔG via PRODIGY^25^ and stability via GROMACS^26^ 2025.4 MD + gmx_MMPBSA,^27^ benchmarked to 4HTC. Of this plan, only a Boltz-2 pilot against thrombin and its calibration were carried out; the full target panel, the AlphaFold3-multimer arm, PRODIGY, MD and MM-GBSA were not run. The pilot is reported here rather than omitted, because withholding an executed but unsupportive experiment would be a selective omission.

In the pilot, the hirudin×thrombin positive control reached ipTM 0.6613 and the candidate g21166×thrombin 0.6165, against a background of 20 independent public decoys (0/20 and 1/20 decoys reached or exceeded them, respectively). Repeat runs of the candidate under fixed seeds were seed-sensitive and insufficiently reproducible (range 0.3346 across four repeats, larger than its 0.2506 separation from the decoy mean), whereas the positive control varied far less; with n = 4 this describes reproducibility, and we do not interpret it as a property of the candidate itself.

A separate comparison panel of 28 objects, run under the same input conditions and seed design (referred to below as the null panel; its members are not verified non-binders) — 20 independent public decoys plus 8 non-lead cargo proteins from this study — run at 3 input conditions × 10 seeds (28 × 3 × 10 = 840 predictions; the candidate and the positive control were run under the same design as a further 2 × 3 × 10 = 60, 900 in total, and are not members of the 28) showed that supplying an MSA raises ipTM for decoys as well (15/20 decoys increased; median +0.1029) and that the positive control’s increase (+0.1406) sat at the 65th percentile of that decoy distribution — i.e. under this design the MSA-driven ipTM gain is not discriminative. That is a statement about the assay’s resolving power, not evidence that the candidate does not bind. In the same 840 predictions we did not observe a consistent improvement in *reproducibility* either — improvement in some objects and deterioration in others can produce a near-zero overall median, so this is an absence of a consistent panel-level effect, not evidence that MSAs do not affect reproducibility: across the 28 comparison objects the seed-to-seed ipTM range narrowed in 15 and widened in 13 (paired median change -0.014), and the interquartile range narrowed in 14 and widened in 14 (median -0.001).

Seed-to-seed stability nevertheless separates the two reference points completely within this fixed panel: under unpaired MSA the positive control’s seed range (0.0810) was smaller than that of every one of the 28 comparison objects, and g21166’s (0.6362) was larger than that of every one of the 28. Adding each of them in turn to those 28 gives a 29-object set in which the positive control ranks first by ascending range and g21166 ranks first by descending range; both are descriptive ranks. We report these as ranks rather than as *p* values, because with a fixed challenge panel the corresponding tail fraction (1/29) is an empirical rank under an exchangeability assumption, not a test against a defined null population. Level and stability could in principle be confounded, since a score near the ceiling has less room to vary; within the 28 comparison objects the observed monotonic association was weak (Spearman rho = -0.17; median range 0.335 in the upper half of the level distribution versus 0.361 in the lower half), but this is a descriptive comparison without a confidence interval and does not establish independence, and it spans only the comparison objects’ ipTM range (0.25-0.73), not the positive control’s 0.81, so a ceiling effect at the positive end cannot be excluded. With a single positive control these observations cannot show that low seed variance distinguishes binders from non-binders. Nor are they a calibration of the n = 4 pilot statement above: a range grows mechanically with the number of repeats, so an n = 4 range and an n = 10 range are not the same estimator. They provide an external reference at a larger sampling scale for the same phenomenon -seed sensitivity - and no more.

For completeness, in the paired-versus-unpaired MSA contrast the candidate’s gain (+0.1419) was larger than that of every one of the 20 public decoys (the candidate is not itself a decoy); this is an uncalibrated descriptive percentile, is confounded by MSA depth (decoy depths span 2–2505), and the group-level contrast showed no effect (12 of 20 decoys increased; median +0.0036), so it is recorded as an exploratory observation and no claim is built on it. Accordingly this paper makes no positive target-engagement claim for any candidate.

### Offline security

No candidate sequence, structure or alignment was submitted to any online prediction service; all folding/search ran locally, with only public reference data downloaded inbound.

## Results and Discussion

### 2.1 Starting point: the ultra-enriched candidate set and its interactome context

We take as input the 17 ultra-enriched candidate cargo proteins nominated by the companion resource^1^ (log2FC ≥ 10, P < 0.05; almost exclusively detected in the salivary-EV fraction, with bootstrap-CI-stabilised fold-changes reported in ref 1). We do not re-derive the differential-abundance analysis here; the compartment structure, EV signature and pharmacopoeia isoform divergence are established in ref 1 and are not reproduced here.

One new, orthogonal observation motivates the structural focus. In the network represented by the node and edge tables used here (507 proteins, giant component 387; the plotting script records these as a STRING network filtered at score ≥900, an attribute we have not independently verified — see Experimental Section), the bulk cargo resolves into a powder-derived mitochondrial/ribosomal core and salivary-EV membrane-trafficking clusters — but the disulfide-rich dark leads (g21166, g436, g20791) and g8114 — the secreted protein carrying predicted Trypsin-like segments — do not appear among the nodes of the network tables queried (the original orthology-mapping and interaction-filtering records are not available to us, so we do not describe their outcome; Figure 1). This non-appearance does not demonstrate a lack of function; it is what motivates the structural analysis that follows.

**Figure 1.**
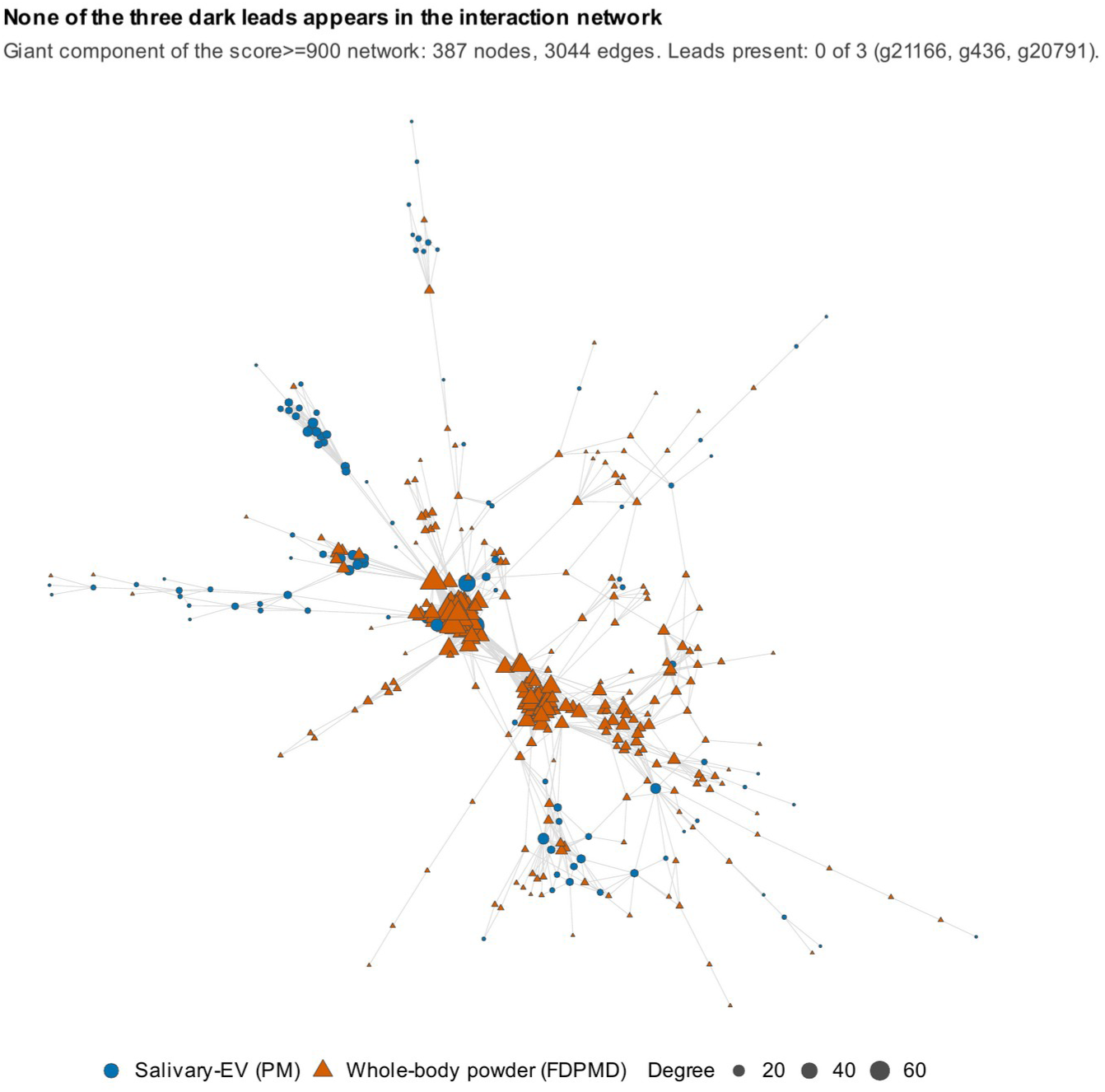
Candidate interactome context. The three lead candidates do not appear among the nodes of the network tables used here.

### 2.2 A small, cysteine-dense “dark” fraction resists sequence-based annotation

Physicochemical and disulfide characterization of the small candidates separates them into two classes. The high-abundance cargo comprises intracellular/membrane machinery with clear Pfam domains (Ras/Arf GTPases, calcium-transporting ATPases, calpain [Peptidase_C2], voltage-gated Cl⁻ channel, RING fingers) — consistent with the membrane/GTPase cargo of a vesicle-secreted compartment. In sharp contrast, three candidates form a small, secreted, cysteine-dense group with no Pfam domain at E<1e-5 (Table 1).

**Table 1.** Sequence features of the three dark-fraction candidates. Pfam, hmmscan against Pfam-A at E < 10⁻⁵; putative S–S, the maximum number of disulfide bonds allowed by the cysteine count (⌊Cys/2⌋).

| **Candidate** | **Length** | **Cys** | **Putative S–S** | **Pfam** | **Note** |
|---|---|---|---|---|---|
| g21166.t1 | 135 aa | 10 | 5 | none | Protein-sleepless homolog<sup>1</sup>; 10-Cys pattern consistent with a Ly-6/uPAR three-finger fold |
| g436.t1 | 171 aa | 12 | 6 | none | uncharacterised; protease-inhibitor-like disulfide density |
| g20791.t1 | 147 aa | 4 | 2 | none | uncharacterised |

Separately, g8114.t1 carries Pfam Trypsin (PF00089) serine-protease sequence in three domain envelopes (i-Evalue 3.6e-20, 2.8e-15 and 3.3e-06 at residues 156–354, 633–728 and 1176–1258; a fourth envelope at 1250–1311 has i-Evalue 0.35 and does not meet the E < 1e-5 cutoff used here) together with LDL-receptor-A and F5/8-type-C modules — a secreted, multidomain serine protease of undetermined specificity; the two envelopes carrying the catalytic residues (633–728 and 1176–1258) are the subject of Section 2.4. The absence of a Pfam assignment for the dark trio, despite bona-fide secretion signals and high disulfide density, is precisely why the companion study flagged them for structural follow-up.

### 2.3 Predicted structures assign the flagship candidate to the Ly-6/uPAR three-finger fold

We folded the six small (≤260 aa) ultra-enriched candidates with ESMFold (single-sequence, CPU). Model confidence varied (mean pLDDT 0.30–0.80); the disulfide-densest dark leads folded at lower confidence (g436 0.40; g20791 0.30) — an expected limitation of single-sequence prediction for cysteine-rich micro-proteins and the reason MSA-based modelling is scoped as the extension (Section 2.5, Discussion). Pairwise TM-align against a panel of canonical anticoagulant/toxin folds (Kunitz, Kazal, potato-inhibitor-I/eglin, hirudin, antistasin/hirustasin, three-finger/Ly-6) gave a decisive result for the flagship lead:

- the predicted structure of g21166.t1 supports assignment to the Ly-6/uPAR three-finger fold (TM-align normalized by the reference domain: 0.58 vs human CD59, 0.56 vs α-bungarotoxin; same-fold threshold TM≥0.5), with a chemically sensible set of five predicted disulfide pairings (Cys29–60, 32–41, 55–77, 87–100, 102–108; predicted, not measured) matching the ∼4–5 disulfides of the three-finger scaffold. Whereas the companion study^1^ nominated g21166 as a Protein-sleepless / Ly-6/uPAR homolog on sequence grounds, the predicted structure supports that fold assignment — assigning the leech Protein-sleepless homolog to the Ly-6/uPAR three-finger superfamily. We explicitly do not infer a function from this assignment: the same fold carries complement regulation (CD59), protease-receptor (uPAR), cell-adhesion and nicotinic-modulator (Lynx1/SLURP) activities, and experimental work on this superfamily shows that fold-level similarity does not transfer function. A Foldseek search against AFDB-SwissProt (∼540k reviewed AlphaFold structures; search products deposited with this study) returns 23 significant matches for g21166 (alnTM ≥ 0.5 and E ≤ 0.05), the strongest being a U-scoloptoxin at alnTM 0.630 (E = 4.4×10⁻⁴) and a three-finger toxin (MALT0057C) at alnTM 0.587 (E = 4.5×10⁻3). Note that its single highest-alnTM hit (alnTM 0.672) is *not* significant (E = 0.12) and is therefore not used here. Because `alntmscore` is normalized by min(alignment, query, target) length, these matches corroborate the assignment made by full-length TM-align above but do not by themselves constitute a same-fold assignment. The PDB100 search — a different database source, not an independent algorithm or query — reinforces the assignment directly: g21166’s best significant PDB match is Lynx1 (PDB 2L03, water-soluble Lynx1; alnTM 0.664, E = 0.017), with two further significant matches to C4.4A (6ION/6IOM) — both members of the Ly-6/uPAR three-finger family to which we assign it, and Lynx1 itself the prototypical nicotinic-acetylcholine-receptor modulator.
- The remaining dark leads matched no panel fold above threshold (g436 best TM 0.38 vs eglin; g20791 0.29 vs Kazal). Because their models are low-confidence (pLDDT 0.30–0.40), we report these as not confidently assignable rather than claiming novel folds; MSA-based modelling (ColabFold/AlphaFold3, offline) is the explicit next step. Both TM normalizations are included in the reproducibility archive described under Data Availability.

Structurally, the g21166 match is carried by the disulfide-bonded core (res 29–108), which superposes on CD59 and α-bungarotoxin (cealign RMSD 4.7 Å/56 res vs bungarotoxin; 6.4 Å/64 res vs CD59) with the three β-hairpin “fingers” and the five predicted disulfide pairings recapitulated (Figure 2); the model’s lower-confidence N/C-terminal segments are excluded from the fold call. g21166 is thus a three-finger-domain-containing protein, not necessarily a pure three-finger toxin.

**Figure 2.**
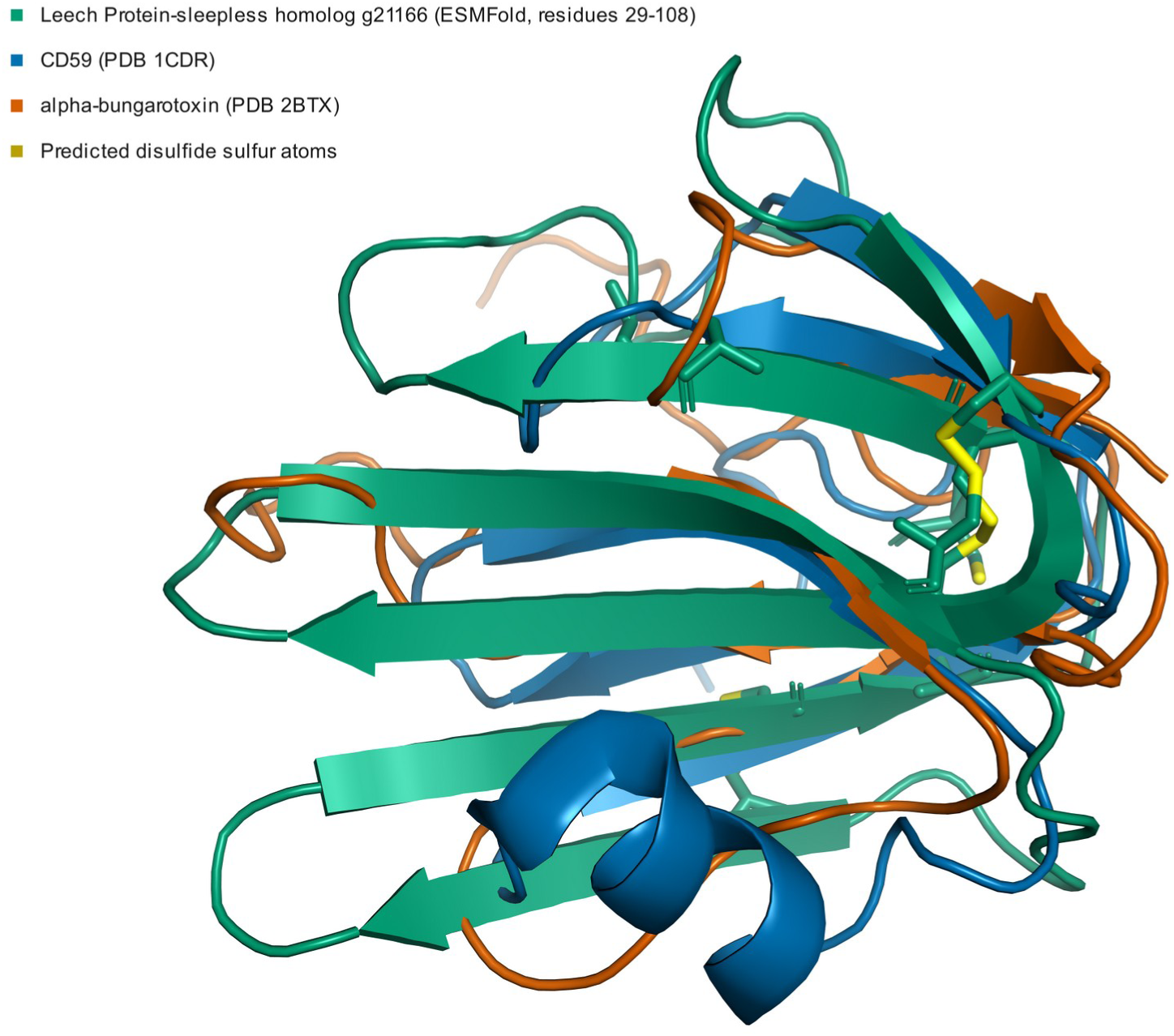
Predicted three-finger fold of g21166. Predicted model of the leech Protein-sleepless homolog superposed on Ly-6/uPAR three-finger references; five predicted disulfide pairings are indicated.

### 2.4 Structural-search status of the dark fraction

Beyond the three-finger assignment, the two remaining dark scaffolds behave differently from one another once all 16 modelled candidates (of 17 nominated; Table S1) are searched against AFDB-SwissProt (2026-08-01 run, foldseek 10.941cd33, same parameters as the earlier search): g20791 has no Foldseek hit with alnTM ≥ 0.5 (maximum alnTM 0.222; and in the PDB100 search only a single hit at all, alnTM 0.178), whereas g436 has exactly one hit above the TM threshold (alnTM 0.596, a nuclear RNA-polymerase subunit) and that hit is not statistically significant (E = 9.5), so by the criterion above neither g20791 nor g436 has a significant Foldseek match — in contrast to g21166, which has 23. The PDB100 search likewise gives the same criterion-level result: g436 and g20791 again return zero significant matches (g436’s highest is alnTM 0.613 at E = 2.6), whereas g21166 returns five. No match meeting the prespecified criterion was detected for either candidate in the two database searches. This concordant no-match status is not evidence of a novel fold: it remains conditional on query-model confidence — the ESMFold models for g436 and g20791 have mean pLDDT of only 0.4 and 0.3 respectively — as well as on the database snapshots and the Foldseek settings stated in the Experimental Section, and both searches used the same algorithm on the same low-confidence queries.

We therefore tested that confidence limitation directly for g436. A local MMseqs2 search of the 121,389,642-entry UniRef90 snapshot returned 102 homologous sequences for g436 (A3M depth 103) but no additional sequence at all for g20791 (query-only A3M, depth 1; a result reproduced byte-for-byte in two independent runs). g436 is therefore sequence-visible: homologous sequence information exists in this database, although this does not by itself explain the low confidence of the original ESMFold model. In a within-model ablation (same Boltz-2 setting, same parameters, three seeds per arm, only the MSA input differing), supplying the depth-103 MSA increased mean complex pLDDT from 0.471 to 0.709. The increase was localized to the homolog-covered mature region (residues 49–171; mean residue-wise pLDDT 0.440 without MSA versus 0.781 with MSA), whereas residues 1–48 showed no improvement (0.550 versus 0.524). Foldseek searches of the highest-confidence MSA-based prediction nevertheless detected no match meeting the prespecified joint criterion (alnTM ≥ 0.5 and E ≤ 0.05) in either PDB100 or AFDB-SwissProt. The no-match status therefore cannot be explained solely by the low confidence of the original single-sequence query model; this does not establish a novel fold and remains conditional on prediction accuracy, database snapshots, and the Foldseek settings stated in the Experimental Section. Although the no-MSA arm produced higher maximum alnTM values (0.848/0.865), none satisfied the E-value component of the joint criterion, illustrating why alnTM alone was not used for classification. Neither, therefore, returned a match to a canonical anticoagulant or toxin fold that meets the prespecified criterion in the databases, models and settings used — a non-detection, not a demonstrated absence.

The secreted protein g8114 carries two predicted Trypsin-like segments; a complete catalytic unit, and any activity, are not established here. The motifs that correspond to the conserved catalytic histidine (TAAHC, residue 670) and catalytic serine (GDSGGP, residue 1239) lie 569 residues apart, in two separate Pfam Trypsin segments (633–728 and 1176–1258), with F5/8 type C, LDL-receptor class A and SRCR domains interposed between them. Its closest AFDB-SwissProt neighbour by alignment-normalised TM is the snake-venom serine protease Achelase-2 (alnTM 0.882), but that alignment spans only residues 642–714 — 73 of 1313 (5.6% of the query; TM normalised by query length 0.054) and falls inside the histidine-containing segment — it is a local match, not whole-protein similarity. Two lines of evidence support conserved genomic organisation of this region and splicing of part of it, but neither establishes a complete 1,313-aa transcript or an active catalytic unit, and neither excludes a merged gene model: First, protein-to-genome alignment of the 1,313-aa model onto three chromosome-level hirudinean assemblies recovers a single collinear locus in each (*H. manillensis* chr10, 32 exons, identity 1.000; *Hirudo nipponia* chr8, 30 exons, 0.523; *Whitmania pigra* chr7, 30 exons, 0.513), with intron positions conserved at a ±3-aa tolerance (28/31 and 29/31, including 12/12 and 11/12 of the introns that fall between the two Trypsin segments). The reading frame is frameshift-free and 3,942 bp only in *H. manillensis*, the assembly from which the model derives; the two outgroup alignments each carry frameshifts and one internal stop codon (3 and 1 in *H. nipponia*, 2 and 1 in *W. pigra*), as expected at ∼0.52 protein identity, so they support conservation of exon–intron architecture but not an intact outgroup reading frame. Because one query protein and one aligner were used for all three genomes, these three mappings are not independent of one another and on their own test conservation of exon–intron architecture rather than transcript identity. Second, and independently of the query protein, splice junctions extracted from public *H. manillensis* head RNA-seq show that the intervening region is spliced in mature transcript: 11 of its 12 predicted introns carry junction-spanning reads in each of two libraries from two separate studies (SRR26151944, 42.8× regional depth; SRR26541753, 6.3×), with no evidence of a lower detection rate than expression-matched control introns on the same chromosome (one-sided lower-tail binomial *P* = 0.902 and 0.984; this is a non-rejection, not an equivalence test, and no equivalence margin was defined), and the second Trypsin segment is spliced as well (1/2 introns, *P* = 0.341 on the merged head data at 42.5×). The same test on salivary-gland libraries detects none of these 12 junctions (0/12, *P* = 4.3 × 10⁻⁴), so the junctions are detected in the head libraries and not detected in the salivary-gland libraries; because the two tissue sets come from different studies, a tissue effect cannot be separated from a study/batch effect, and we therefore do not attribute the difference to tissue. Two further head libraries were excluded by a prespecified read-count rule before analysis, so two libraries carry the positive result. The C-terminal segment beyond residue 1258 remains undetermined at the transcript level. No de-novo genome annotation was performed; the analysis used protein-to-genome alignment against published assemblies and read alignment to the published *H. manillensis* assembly (see Experimental Section). The dark fraction therefore contains at least one confidently-assigned three-finger scaffold and two structurally unresolved cysteine-rich micro-proteins whose no-match status is the primary target of the MSA-based structural extension. (Figure 3; the g436/g20791 no-match status is conditional on the models, databases and thresholds stated.)

**Figure 3.**
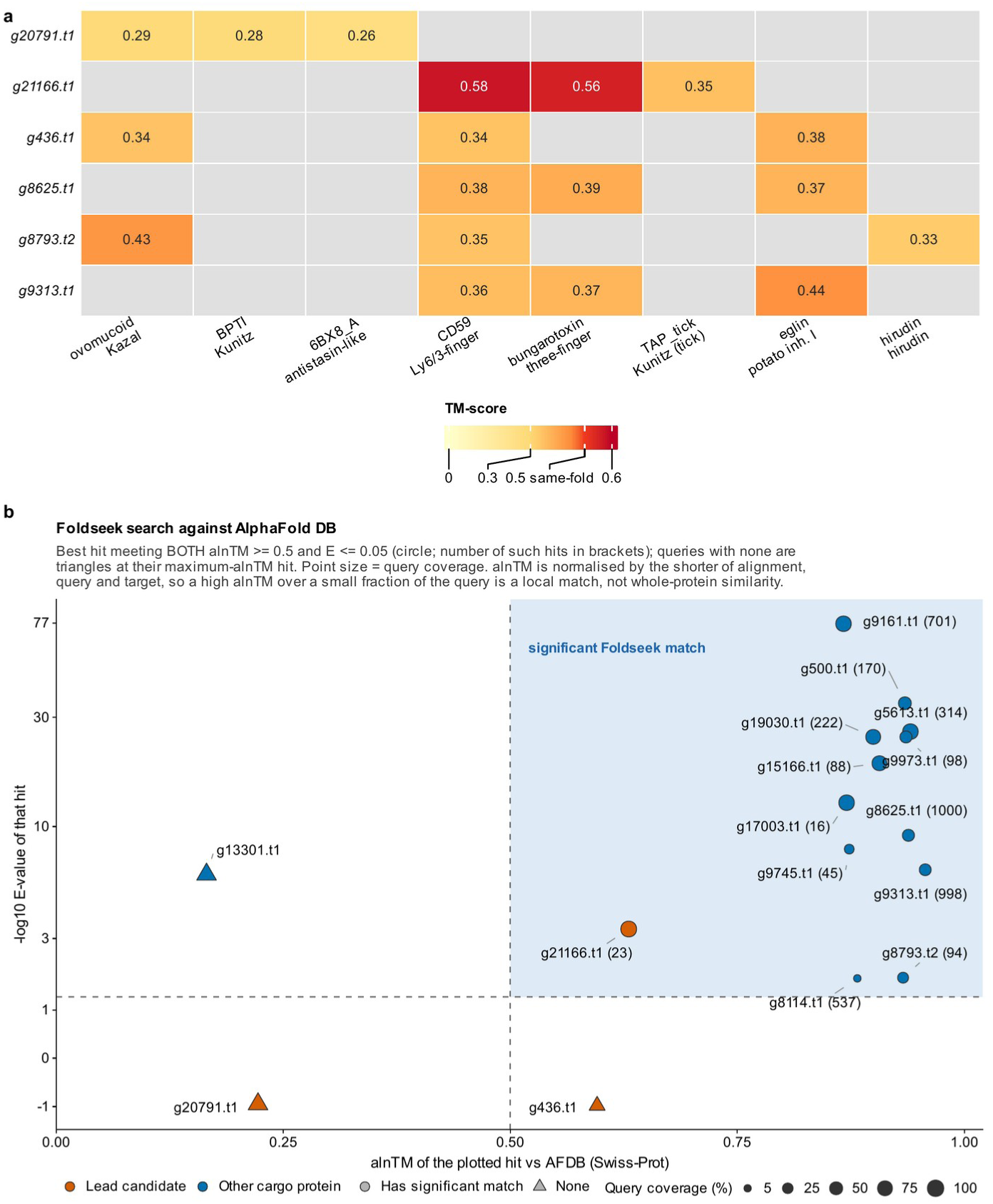
Structural-search status of the dark fraction. TM-analog matrix (candidate × reference) together with Foldseek search status. Of the 60 candidate×reference pairs tested, 2 reach the same-fold threshold, both for g21166 against three-finger references (TM 0.58 and 0.56); the heatmap shows the top three references per candidate. In the AFDB-SwissProt search, evaluated over all 8,665 reported hits, a *significant Foldseek match* (alnTM ≥ 0.5 and E ≤ 0.05) exists for 12 of the 13 non-lead cargo proteins searched and for 1 of the 3 leads: g21166 has 23 such matches (best alnTM 0.630, E 4.4×10⁻⁴, U-scoloptoxin; also three-finger toxin MALT0057C at 0.587), whereas g436 and g20791 have none.

### 2.5 Target engagement: an unsupportive thrombin pilot and a terminated nicotinic test

The structural assignments define the functional tests. g21166 is assigned to the Ly-6/uPAR fold, which does not by itself nominate a target; g8114 is a secreted protein carrying Trypsin-like segments whose substrate specificity is unknown (no fibrino(geno)lytic activity is claimed or tested here); and coagulation-factor inhibitor engagement (thrombin/FXa/FXIa) is most meaningfully tested on the classical pharmacopoeia protease-inhibitor families. The target-engagement layer was designed as the core intended contribution of this paper: high-confidence (MSA/AlphaFold3) models of each lead, AlphaFold3-multimer/Boltz-2 complex prediction with the human target panel, and PRODIGY ΔG and short GROMACS MD + MM-GBSA re-scoring benchmarked against the experimental hirudin–thrombin interface (4HTC). Of this design, only a Boltz-2 thrombin pilot was executed (Experimental Section), and it does not support a positive target-engagement claim. Although the compute prerequisites were met, the remaining complexes were not modelled, for methodological rather than computational reasons, and the pre-registered nicotinic interface test was terminated as undecidable at the null-model construction stage (Experimental Section; Supplementary Note S1). We make no functional-activity claim. Comparison-set members are described as having no experimental evidence of interaction with the nicotinic target; they are not verified non-binders, and no statement in this work should be read as establishing that any of them fails to bind.

### 2.6 A prioritized, structure-informed candidate shortlist

Integrating secretion features, disulfide density, domain status and fold assignment (Table 2) stratifies the leads into three tiers. Priority-1: three small, secreted, cysteine-rich leads — g21166.t1 (Ly-6/uPAR three-finger fold; the best-supported fold assignment in the set) and the two disulfide-dense “dark” scaffolds g436.t1 and g20791.t1. Priority-2: the secreted protein g8114.t1, which carries predicted Trypsin-like segments of unknown specificity. Priority-3: the abundant but intracellular/membrane cargo, which reflects vesicular-packaging biology rather than direct therapeutic leads. This structure-informed tiering — not raw abundance — is the actionable output (the single most-enriched protein, a Ras-family GTPase, is Priority-3) (Table 2 and Figure 4).

**Table 2.** Structure-informed prioritization of the 17 candidate cargo proteins. Priority 1, small secreted cysteine-rich leads; 2, secreted protein carrying predicted Trypsin-like segments; 3, abundant intracellular/membrane cargo. Closest panel fold, best pairwise TM-align match against the reference fold panel (TM normalized by the reference domain); a fold is assigned only at TM ≥ 0.5. The panel comparison and mean pLDDT refer to the single-sequence ESMFold models of the six candidates ≤ 260 aa; n.a., not assessed. log2FC, salivary-EV-enriched versus whole-body fraction, from ref 1.

| Candidate | Priority | Annotation | Length (aa) | Cys | Putative S–S | Closest panel fold (TM) | Mean pLDDT | log2FC |
|---|---|---|---|---|---|---|---|---|
| g436.t1 | 1 | Uncharacterized protein | 171 | 12 | 6 | none (best potato inhibitor I, 0.38) | 0.4 | 13.08 |
| g21166.t1 | 1 | Protein sleepless | 135 | 10 | 5 | Ly-6/uPAR three-finger (0.58) | 0.6 | 11.23 |
| g20791.t1 | 1 | Uncharacterized protein | 147 | 4 | 2 | none (best Kazal, 0.29) | 0.3 | 10.52 |
| g8114.t1 | 2 | Peptidase S1 domain-containing protein | 1313 | 68 | 34 | n.a. | n.a. | 10.05 |
| g8625.t1 | 3 | Uncharacterized protein | 260 | 5 | 2 | none (best three-finger, 0.39) | 0.8 | 15.78 |
| g9161.t1 | 3 | Calcium-transporting ATPase | 1018 | 16 | 8 | n.a. | n.a. | 13.02 |
| g13301.t1 | 3 | Phosphatase and actin regulator 1 | 525 | 9 | 4 | n.a. | n.a. | 11.84 |
| g15166.t1 | 3 | Phosphatidic acid phosphatase type 2/haloperoxidase domain-containing protein | 340 | 11 | 5 | n.a. | n.a. | 11.17 |
| g5613.t1 | 3 | Sodium/solute symporter | 610 | 18 | 9 | n.a. | n.a. | 11.09 |
| g8793.t2 | 3 | RING-type domain-containing protein | 222 | 20 | 10 | none (best Kazal, 0.43) | 0.6 | 11.07 |
| g500.t1 | 3 | Calpain catalytic domain-containing protein | 664 | 13 | 6 | n.a. | n.a. | 10.91 |
| g9973.t1 | 3 | Phospholipid-transporting ATPase | 578 | 12 | 6 | n.a. | n.a. | 10.73 |
| g18761.t1 | 3 | HEPN domain-containing protein | 3570 | 69 | 34 | n.a. | n.a. | 10.65 |
| g9313.t1 | 3 | Cell division control protein 42 homolog | 209 | 4 | 2 | none (best potato inhibitor I, 0.44) | 0.8 | 10.45 |
| g9745.t1 | 3 | Cytochrome b561 domain-containing protein | 631 | 11 | 5 | n.a. | n.a. | 10.19 |
| g19030.t1 | 3 | SLC26A/SulP transporter domain-containing protein | 699 | 6 | 3 | n.a. | n.a. | 10.13 |
| g17003.t1 | 3 | Uncharacterized protein | 337 | 14 | 7 | n.a. | n.a. | 10.10 |

**Figure 4.**
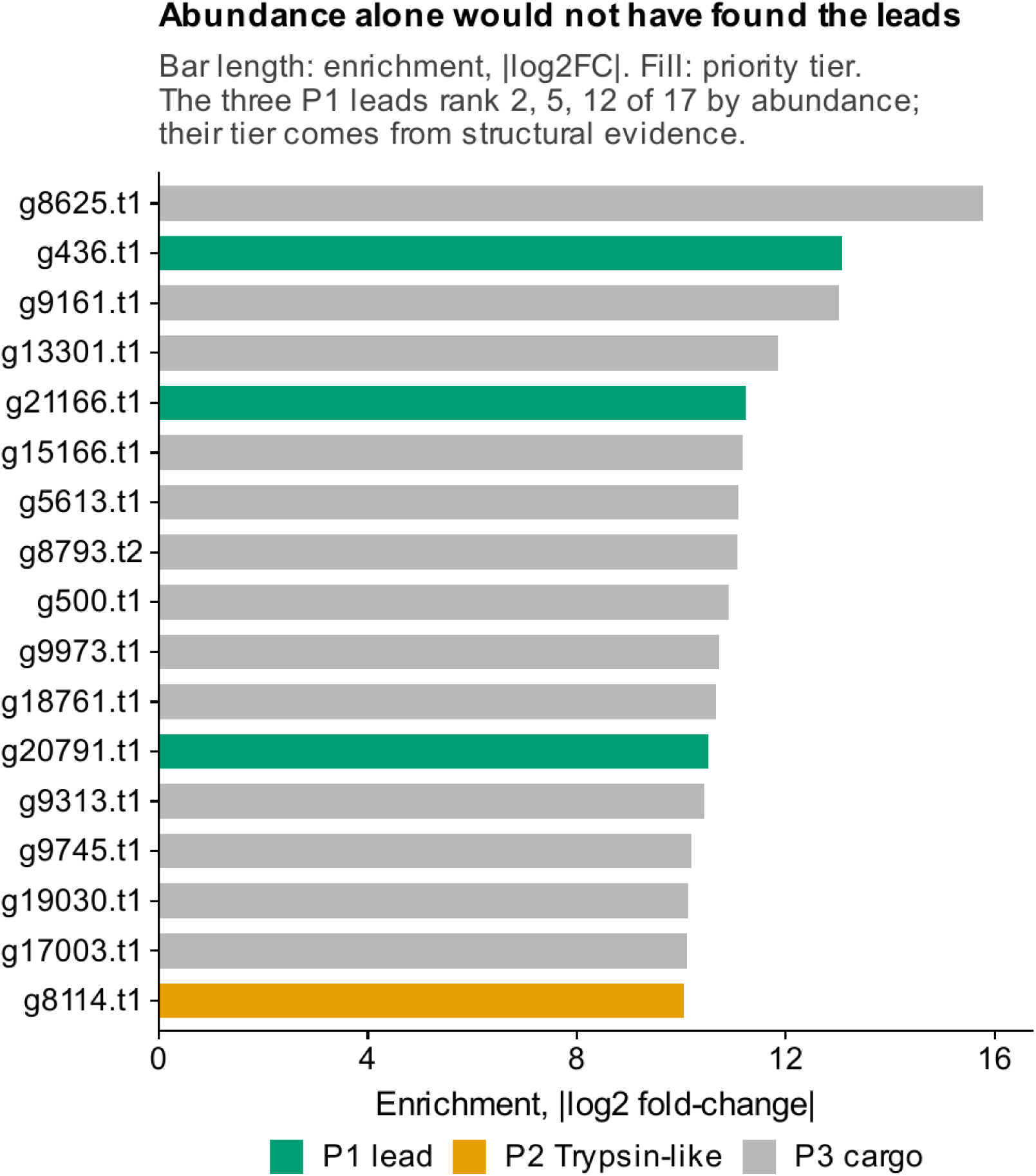
Prioritized structure-informed shortlist. Bars show the enrichment of each of the 17 candidates (|log2 fold-change|, salivary-EV-enriched versus whole-body fraction, from ref 1); fill shows the structure-informed priority tier (P1, small secreted cysteine-rich leads; P2, secreted protein carrying predicted Trypsin-like segments; P3, intracellular/membrane cargo). The three P1 leads rank 2, 5 and 12 of 17 by enrichment, so the tiers derive from structural evidence rather than abundance.

### 2.7 Discussion

This study asked whether structure-based annotation can resolve salivary-EV cargo that sequence methods leave uninterpretable. For one of the three sequence-dark candidates it can: the leech Protein-sleepless homolog g21166 is assigned to the Ly-6/uPAR three-finger fold by full-length TM-align against two references, by 23 significant AFDB-SwissProt matches and by a significant PDB match to Lynx1, converting a sequence-level nomination^1^ into a fold-level hypothesis. Three-finger proteins have been described in invertebrates such as starfish,^5^ and g21166 adds a candidate from a hematophagous annelid, packaged in salivary EVs, to that comparative picture. We infer no function from the fold: the Ly-6/uPAR superfamily spans complement regulation, protease-receptor, cell-adhesion and nicotinic-modulator activities, and no target-engagement result in this work supports a mechanism.

The other two dark candidates show that a missing structural match can have different causes, and the difference matters for what to do next. g20791 has no detectable homolog in a 121-million-entry UniRef90 snapshot and only a low-confidence model, so its status is limited by sequence information. g436, in contrast, has 102 homologs; supplying them raised model confidence (mean complex pLDDT 0.471 to 0.709), yet the homolog-informed model still matched nothing under the prespecified criterion. Neither result demonstrates a novel fold, but g436 is the more informative target for experimental structure determination, because its no-match status cannot be attributed solely to missing homolog information. Together with the absence of the leads from the nodes of the interaction network used here, these results place the least-annotated secreted cargo, rather than the most abundant, at the top of the experimental shortlist (Table 2).

Methodologically, the workflow is offline and criterion-based: a fold is assigned only at full-length TM ≥ 0.5, a structural match requires alnTM ≥ 0.5 and E ≤ 0.05 over all reported hits, and a no-match result is qualified by model confidence and homolog depth. It can therefore be applied to unpublished proteomes that cannot be sent to online services.

### Limitations

All structural and functional statements are predictions with quantified confidence, not measured activities. Single-sequence ESMFold yielded low confidence for the cysteine-densest micro-proteins (pLDDT 0.30–0.40 for g436/g20791), so their no-match status must be interpreted cautiously pending higher-confidence MSA-based (AlphaFold3) modelling. The target-engagement layer that would raise these fold annotations to mechanism-level leads was only partly executed (Section 2.5); the structural content is therefore a fold-annotation advance over ref 1 rather than a functional discovery, and the two papers are designed to be read together. No vesicle authentication (TEM/NTA/Western) was performed; EV-biology statements rest on ref 1, which is cited but not re-claimed here.

## Conclusions

Structure-based annotation resolves part of the sequence-dark salivary-EV cargo of *H. manillensis*. The leech Protein-sleepless homolog g21166 is assigned to the Ly-6/uPAR three-finger fold at the level of prediction, without functional inference. g436 and g20791 remain structurally unresolved for different reasons — g436 despite homolog-informed modelling, g20791 for lack of any detectable homolog — and neither is shown to be a novel fold. The two-Trypsin-segment model g8114 is supported as a conserved locus whose intervening region is spliced in head transcripts, but neither a complete transcript nor a catalytic unit is established, and no target-engagement claim is made. The structure-informed shortlist and the offline, criterion-based workflow provide a starting point for experimental structure determination and activity testing of the leech salivary-EV cargo.

## Data Availability

The mass spectrometry proteomics data underlying this study have been deposited to the ProteomeXchange Consortium via the iProX partner repository with the dataset identifier PXD081595 (iProX accession IPX0018563000); the deposition is shared with the companion study^1^ and will be released publicly upon publication.

The public data reused in this study are the genome assemblies GCA_034509925.1, GCA_040113095.1 and GCA_041430665.1 and the RNA-seq runs of BioProjects PRJNA861860, PRJNA1019887 (SRR26151944) and PRJNA1032729 (SRR26541746, SRR26541752 and SRR26541753); no new sequence data were generated.

The structural products of this study — the 16 predicted models with their run records, the raw Foldseek outputs against AFDB-SwissProt and PDB100, the TM-align fold-assignment matrix, the per-seed Boltz-2 ipTM tables and the g436 MSA-ablation models— will be deposited, together with the network node and edge tables and the analysis scripts, in a Zenodo reproducibility archive upon journal publication; until then they are available from the corresponding author on reasonable request.

## Supporting Information

The following Supporting Information accompanies this preprint:

Supplementary Note S1: rationale for terminating the pre-registered nicotinic null model (PDF)

Table S1: per-candidate model provenance, 17 candidates (CSV) Table 2 source data: prioritized candidate shortlist (CSV)

## Author Contributions

Hongfen Cen, Fang Zhao and Juan Zhao contributed equally to this work. Hongfen Cen: conceptualization, investigation, formal analysis, visualization, writing – original draft. Fang Zhao: investigation, formal analysis, writing – review & editing. Juan Zhao: investigation. Xiangrong Tong: formal analysis. Zichao Liu: formal analysis, writing – review & editing. Feng Zhao: conceptualization, formal analysis, supervision, writing – review & editing. All authors have read and approved the submitted version.

## Acknowledgments

The authors thank the funding bodies listed under Funding.

## References

(1) Zhao, F.; Zhao, J.; Tong, X.-R.; Wang, M.; Liu, Z.; Zhao, F. A Comparative Astral DIA Proteome of the Asian Medicinal Leech *Hirudinaria manillensis*: Salivary EV-Enriched Fraction versus Whole-Body Pharmacopoeial Powder. PeerJ, submitted (ms 144559), 2026.

(2) Jumper, J.; Evans, R.; Pritzel, A.; Green, T.; Figurnov, M.; Ronneberger, O.; Tunyasuvunakool, K.; Bates, R.; Žídek, A.; Potapenko, A.;, et al. Highly accurate protein structure prediction with AlphaFold. Nature 2021, 596 (7873), 583–589. DOI: 10.1038/s41586-021-03819-2.

(3) van Kempen, M.; Kim, S. S.; Tumescheit, C.; Mirdita, M.; Lee, J.; Gilchrist, C. L. M.; Söding, J.; Steinegger, M. Fast and accurate protein structure search with Foldseek. Nat Biotechnol 2024, 42 (2), 243–246. DOI: 10.1038/s41587-023-01773-0.

(4) Koh, K.; Joiner, W. J.; Wu, M. N.; Yue, Z.; Smith, C. J.; Sehgal, A. Identification of SLEEPLESS, a sleep-promoting factor. Science 2008, 321 (5887), 372–376. DOI: 10.1126/science.1155942.

(5) Lyukmanova, E. N.; Bychkov, M. L.; Chernikov, A. M.; Kukushkin, I. D.; Kulbatskii, D. S.; Shabelnikov, S. V.; Shulepko, M. A.; Zhao, R.; Guo, W.; Kirpichnikov, M. P.;, et al. In Search of the Role of Three-Finger Starfish Proteins. Mar Drugs 2024, 22 (11), 488. DOI: 10.3390/md22110488.

(6) Welsh, J. A.; Goberdhan, D. C. I.; O’Driscoll, L.; Buzas, E. I.; Blenkiron, C.; Bussolati, B.; Cai, H.; Di Vizio, D.; Driedonks, T. A. P.; Erdbrügger, U.;, et al. Minimal information for studies of extracellular vesicles (MISEV2023): From basic to advanced approaches. J Extracell Vesicles 2024, 13 (2), e12404. DOI: 10.1002/jev2.12404.

(7) Demichev, V.; Messner, C. B.; Vernardis, S. I.; Lilley, K. S.; Ralser, M. DIA-NN: neural networks and interference correction enable deep proteome coverage in high throughput. Nat Methods 2020, 17 (1), 41–44. DOI: 10.1038/s41592-019-0638-x.

(8) Eddy, S. R. Accelerated Profile HMM Searches. PLoS Comput Biol 2011, 7 (10), e1002195. DOI: 10.1371/journal.pcbi.1002195.

(9) Paysan-Lafosse, T.; Andreeva, A.; Blum, M.; Chuguransky, S. R.; Grego, T.; Pinto, B. L.; Salazar, G. A.; Bileschi, M. L.; Llinares-López, F.; Meng-Papaxanthos, L.;, et al. The Pfam protein families database: embracing AI/ML. Nucleic Acids Res 2025, 53 (D1), D523–D534. DOI: 10.1093/nar/gkae997.

(10) Lin, Z.; Akin, H.; Rao, R.; Hie, B.; Zhu, Z.; Lu, W.; Smetanin, N.; Verkuil, R.; Kabeli, O.; Shmueli, Y.;, et al. Evolutionary-scale prediction of atomic-level protein structure with a language model. Science 2023, 379 (6637), 1123–1130. DOI: 10.1126/science.ade2574.

(11) Wolf, T.; Debut, L.; Sanh, V.; Chaumond, J.; Delangue, C.; Moi, A.; Cistac, P.; Rault, T.; Louf, R.; Funtowicz, M.; et al. Transformers: State-of-the-Art Natural Language Processing. In Proceedings of the 2020 Conference on Empirical Methods in Natural Language Processing: System Demonstrations; Association for Computational Linguistics, 2020; pp 38–45. DOI: 10.18653/v1/2020.emnlp-demos.6.

(12) Zhang, Y.; Skolnick, J. TM-align: a protein structure alignment algorithm based on the TM-score. Nucleic Acids Res 2005, 33 (7), 2302–2309. DOI: 10.1093/nar/gki524.

(13) Berman, H. M.; Westbrook, J.; Feng, Z.; Gilliland, G.; Bhat, T. N.; Weissig, H.; Shindyalov, I. N.; Bourne, P. E. The Protein Data Bank. Nucleic Acids Res 2000, 28 (1), 235–242. DOI: 10.1093/nar/28.1.235.

(14) Varadi, M.; Bertoni, D.; Magana, P.; Paramval, U.; Pidruchna, I.; Radhakrishnan, M.; Tsenkov, M.; Nair, S.; Mirdita, M.; Yeo, J.;, et al. AlphaFold Protein Structure Database in 2024: providing structure coverage for over 214 million protein sequences. Nucleic Acids Res 2024, 52 (D1), D368–D375. DOI: 10.1093/nar/gkad1011.

(15) UniProt Consortium. UniProt: the Universal Protein Knowledgebase in 2025. Nucleic Acids Res 2025, 53 (D1), D609–D617. DOI: 10.1093/nar/gkae1010.

(16) Suzek, B. E.; Wang, Y.; Huang, H.; McGarvey, P. B.; Wu, C. H.; UniProt Consortium. UniRef clusters: a comprehensive and scalable alternative for improving sequence similarity searches. Bioinformatics 2015, 31 (6), 926–932. DOI: 10.1093/bioinformatics/btu739.

(17) Steinegger, M.; Söding, J. MMseqs2 enables sensitive protein sequence searching for the analysis of massive data sets. Nat Biotechnol 2017, 35 (11), 1026–1028. DOI: 10.1038/nbt.3988.

(18) Passaro, S.; Corso, G.; Wohlwend, J.; Reveiz, M.; Thaler, S.; Ram Somnath, V.; Getz, N.; Portnoi, T.; Roy, J.; Stark, H.;, et al. Boltz-2: Towards Accurate and Efficient Binding Affinity Prediction. *<u>bioRxiv</u>* 2025. DOI: 10.1101/2025.06.14.659707. Preprint, not peer reviewed.

(19) Homma, F.; Lyu, J.; van der Hoorn, R. A. L. Using AlphaFold Multimer to discover interkingdom protein-protein interactions. Plant J 2024, 120 (1), 19–28. DOI: 10.1111/tpj.16969.

(20) Li, H. Protein-to-genome alignment with miniprot. Bioinformatics 2023, 39 (1), btad014. DOI: 10.1093/bioinformatics/btad014.

(21) Kim, D.; Paggi, J. M.; Park, C.; Bennett, C.; Salzberg, S. L. Graph-based genome alignment and genotyping with HISAT2 and HISAT-genotype. Nat Biotechnol 2019, 37 (8), 907–915. DOI: 10.1038/s41587-019-0201-4.

(22) Csárdi, G.; Nepusz, T. The igraph Software Package for Complex Network Research. InterJournal, Complex Systems 2006, 1695.

(23) Szklarczyk, D.; Nastou, K.; Koutrouli, M.; Kirsch, R.; Mehryary, F.; Hachilif, R.; Hu, D.; Peluso, M. E.; Huang, Q.; Fang, T.;, et al. The STRING database in 2025: protein networks with directionality of regulation. Nucleic Acids Res 2025, 53 (D1), D730–D737. DOI: 10.1093/nar/gkae1113.

(24) Abramson, J.; Adler, J.; Dunger, J.; Evans, R.; Green, T.; Pritzel, A.; Ronneberger, O.; Willmore, L.; Ballard, A. J.; Bambrick, J.;, et al. Accurate structure prediction of biomolecular interactions with AlphaFold 3. Nature 2024, 630 (8016), 493–500. DOI: 10.1038/s41586-024-07487-w.

(25) Xue, L. C.; Rodrigues, J. P.; Kastritis, P. L.; Bonvin, A. M.; Vangone, A. PRODIGY: a web server for predicting the binding affinity of protein-protein complexes. Bioinformatics 2016, 32 (23), 3676–3678. DOI: 10.1093/bioinformatics/btw514.

(26) Abraham, M. J.; Murtola, T.; Schulz, R.; Páll, S.; Smith, J. C.; Hess, B.; Lindahl, E. GROMACS: High Performance Molecular Simulations through Multi-Level Parallelism from Laptops to Supercomputers. SoftwareX 2015, 1–2, 19–25. DOI: 10.1016/j.softx.2015.06.001.

(27) Valdés-Tresanco, M. S.; Valdés-Tresanco, M. E.; Valiente, P. A.; Moreno, E. gmx_MMPBSA: A New Tool to Perform End-State Free Energy Calculations with GROMACS. J Chem Theory Comput 2021, 17 (10), 6281–6291. DOI: 10.1021/acs.jctc.1c00645.

