## Supplementary Note S1 for "Structure-resolved fold annotation of the medicinal-leech salivary-EV cargo: a Ly-6/uPAR three-finger scaffold and a structurally dark fraction"

### Contents

Supplementary Note S1. Rationale for terminating the pre-registered nicotinic null model (this file)

Table S1. Per-candidate model provenance, 17 candidates (separate CSV file)

Table 2 source data: prioritized candidate shortlist (separate CSV file)

### Supplementary Note S1. Rationale for terminating the pre-registered nicotinic null model

*Why presumed rather than verified negatives*

This note documents why the nicotinic target-engagement test reports a comparison set of proteins with no experimental evidence of interaction rather than a set of verified non-binders, and why three alternative criteria were evaluated and rejected.

#### Three rejected criteria

**(i) Annotation-keyword exclusion.** Excluding pool members whose annotation names a nicotinic target is not an accepted method in the benchmarking literature, and it failed repeatedly in practice here: snake three-finger toxin nomenclature does not track target class. Four successive keyword rule sets each admitted at least one nicotinic-receptor ligand into the null pool (*Alpha-neurotoxin NTX-4*; *Lynx1* orthologues; *Cobrotoxin*), because names such as *cobrotoxin*, *FS-2* and *NTX-4* carry no target information.

**(ii) Sequence-identity thresholds.** Excluding pool members above an identity cut-off to known binders is a documented failure mode when the positive set is itself internally divergent. Our two snake-derived positive controls, α-bungarotoxin and α-cobratoxin — same family, same target — share only 50.0% identity under gap-free comparison. A threshold calibrated to separate binders from non-binders cannot be set when the binders themselves are that far apart; the criterion is uncalibrated by construction rather than misapplied.

**(iii) Structural evidence from complexes.** The entire PDB contains only a handful of candidate ligand chains co-deposited with a nicotinic-family receptor, and none matches any member of our pool. A zero hit rate here is an absence of evidence, not evidence of absence, and the criterion has no discriminative power at the scale required.

### What the comparison set therefore is

Following published practice, comparison-set members are reported as having no experimental evidence of interaction with the nicotinic target (cf. Martin et al., *JCIM* 2024). They are not verified non-binders. No statement in this work should be read as establishing that any of them fails to bind.

### A residual bias that cannot be removed by selection

Both candidate null families — snake three-finger toxins and Ly-6/uPAR proteins — are themselves enriched for nicotinic-receptor ligands. The standard justification for treating unlabelled pairs as negatives rests on a low prior rate of true positives together with random sampling (0.325–1.5% of protein pairs interact; Dunham & Ganapathiraju, *Molecules* 2021). Sampling within a ligand-enriched family violates that premise directly, and we are not aware of a published remedy for the case where it is violated. This is one of the reasons the test was terminated rather than reported with a caveat.
